# Human intestinal organoids from spondyloarthritis, Crohn’s disease and healthy controls: an integrated workflow for expansion, biobanking, differentiation, and characterization

**DOI:** 10.64898/2026.11.23.768531

**Authors:** Gerlando Natalello, Ronan Soudy, Claire Le Poupon, Pauline Devisme, Louis Dubois, Alicia Brest, Roula Said-Nahal, Maria Antonietta D’Agostino, Maxime Breban, Thomas Bazin, Claire Cherbuy

## Abstract

Human adult stem cell-derived intestinal organoids (HIOs) are self-organized three-dimensional multicellular structures that recapitulate key architectural and cellular features of the human intestinal epithelium. HIOs retain the donor’s genetic background and provide a model for basic and translational research with possible applications in precision medicine. Here, we present an integrated workflow to generate adult human donor-derived organoids from endoscopic caecal biopsies, expand them, and drive their epithelial differentiation. Following primary culture, the workflow uses two in-house-formulated culture media — an expansion medium to maintain the stem cell lineage, and a differentiation medium to promote epithelial maturation. Both formulations are based on conditioned media and represent tunable media with the potential advantage of lower costs. Using this protocol, we generated HIOs from a donor cohort including conditions potentially linked to a spectrum of intestinal inflammation, namely healthy donors, patients with Crohn’s disease, and patients with spondyloarthritis (SpA) — another incurable disease in which intestinal inflammation is suspected to play a key pathogenic role. We illustrated feasibility across clinically diverse donor groups while targeted gene expression and qualitative imaging supported the acquisition of differentiated epithelial features in HIOs. To our knowledge, this is the first study formally describing the generation of intestinal organoids from SpA patients.

**Graphical abstract:** 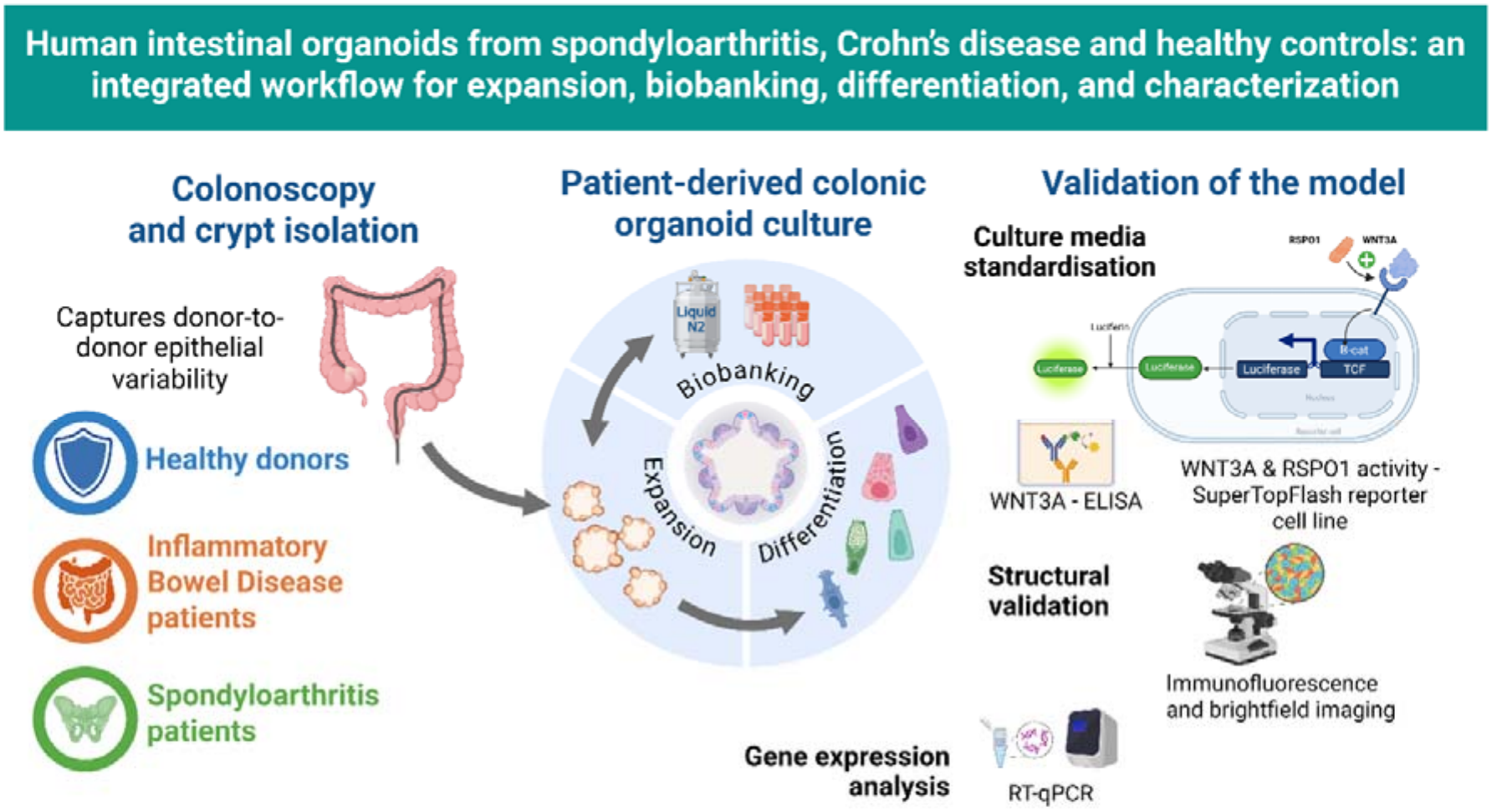

**Highlights:**

- We implemented an integrated workflow to generate and characterize organoids from human caecal biopsies
- The donor cohort spans healthy subjects, spondyloarthritis, and Crohn’s disease patients
- Gene expression and imaging supported differentiated epithelial features in organoids

## INTRODUCTION

Human adult stem cell-derived intestinal organoids (HIOs), obtained from biopsies, are self-organized three-dimensional multicellular structures that reproduce the architecture, cellular diversity, and functions of the human intestinal epithelium [1–3]. HIOs retain the donor’s genetic background and may exhibit a clinically relevant phenotype, making them valuable tools for basic and translational research, as well as precision medicine [4].

These models constitute a complementary alternative to classical preclinical tools — animal models and immortalized cell lines — whose limitations are well documented. Indeed, interspecies physiological and genetic differences may contribute to the high attrition rate of therapies entering clinical trials, including inflammatory bowel disease (IBD) and spondyloarthritis (SpA), while immortalized cell lines, despite their widespread use, lack the cellular complexity needed to reflect human intestinal physiology [5,6]. By recapitulating the diversity and functional organization of the native epithelium, biopsy-derived organoids bridge conventional *in vitro* assays and clinical outcomes, offering a manipulable, patient-specific platform to study disease mechanisms and which may help to predict the toxicokinetics and pharmacokinetics of tested compounds [7]

Given the architectural, cellular, and functional complexity of the human intestinal epithelium — mapped in detail by single-cell analyses [8] — continued efforts to optimize organoid models remain central for building human-relevant platforms of intestinal (patho)physiology. Culture conditions are therefore under continuous refinement to better capture this native cellular complexity: single-cell benchmarking against *in vivo* tissue has been used to identify and correct discrepancies in organoid-derived cell types, such as Paneth cells [9], while more recent protocols have achieved a tunable balance between stem cell self-renewal and differentiation within a single culture system, broadening the cellular diversity captured in human intestinal organoids [10].

Thus, establishing and maintaining robust and relevant models based on HIOs from different groups of donors relies on particularly demanding culture media and conditions [11]. While this sophistication of culture protocols underlies the physiological relevance of organoids, it is also a major source of variability between laboratories. Indeed, despite the exponential growth in publications using gastrointestinal organoids, the reporting of methodologies remains highly heterogeneous across studies, complicating the interpretation, comparison, and reproducibility of results. As highlighted by the recent guiding principles published by VanDussen *et al.* [12], rigorous and standardized reporting of the culture media and conditions used is essential to ensure the reliability and transferability of data generated from gastrointestinal organoid research. This issue is further amplified by the increasing integration of these *in vitro* models into regulatory science and drug development pipelines. HIOs are now recognized as promising components of the Food and Drug Administration’s New Approach Methodologies (NAMs), which aim to reduce reliance on animal testing while improving the predictivity of preclinical assessments [13]. Consequently, ensuring robust standardization, validation, and transparency in organoid culture conditions and reporting is critical to support their qualification for regulatory use and their broader adoption in research [14].

In this work, we present an integrated workflow to generate HIOs from endoscopic caecal biopsies, to expand them, and to drive their epithelial differentiation. This workflow relies on two in-house-produced media — an expansion medium to maintain the stem cell lineage and a differentiation medium to promote epithelial maturation — both formulated using conditioned media, offering a cost-effective alternative to fully recombinant formulations and proprietary commercial media while ensuring compositional transparency and tunability.

Using this protocol, we generated intestinal organoids from a donor cohort capturing a clinically relevant spectrum of gut inflammation, including healthy donors (HCs), patients with overt chronic intestinal inflammation affected by Crohn’s disease (CD), and patients with SpA — another incurable disease in which immuno-inflammation causes a debilitating articular disease [15]. Notably, an altered intestinal homeostasis is suspected to play a key role in SpA based on the frequent presence of subclinical inflammation [16] and proven microbiota [17] and epithelial barrier alterations [18] in the course of the disease. In brief, our protocol is suitable for HIOs generation, characterization and biobanking across three donor groups selected to reflect a clinically relevant gradient of intestinal inflammation: from its absence in HCs, through mild and variable inflammation in SpA [19], to overt inflammation in CD [20], serving as a platform for downstream applications in preclinical research and personalized medicine.

## MATERIAL AND METHODS

### Ethics and human subjects

The study was conducted in accordance with the Declaration of Helsinki and approved by the local Ethics Committee for studies involving humans (reference CNRIPH 21.02.23.68046). Informed consent was obtained from all subjects involved in the study. Participants were enrolled at Hôpital Ambroise-Paré, Assistance Publique–Hôpitaux de Paris (AP-HP), Boulogne-Billancourt, France. Adult individuals were considered for inclusion if they had a clinical indication to undergo colonoscopy. Exclusion criteria were prespecified to ensure participant safety, permit standardized caecal sampling, and minimize biological confounding unrelated to SpA or IBD. Criteria for the definition of each sub-cohort and exclusion criteria are detailed in the Supplementary table 1. A total of 23 subjects were enrolled in the study (8 SpA, 8 HCs, 7 CD).

### Colonoscopy

Colonoscopy was performed according to institutional standard of care using a flexible video endoscope after routine bowel preparation. All biopsies were taken from the caecum, proximal to the ileocaecal valve, from a macroscopically intact area of the mucosa. For each participant, 4 caecal pinch biopsies were collected to provide sufficient material for crypt isolation and parallel analyses. Immediately after removal, biopsies were placed in sterile cold Dulbecco’s Phosphate-Buffered Saline (DPBS, Gibco Ref 14190144) and processed within 4 hours of collection.

### Conditioned media preparation

WNT3A, R-spondin1 (RSPO1), and Noggin (NOG) conditioned media (CM) were produced from available engineered cell lines, all secreting factors cloned from murine transcripts. The L WNT3A cell line was purchased from ATCC (Ref. CRL-2647). HEK293T cells transfected to express murine RSPO1 (HEK-RSPO1) were purchased from Trevigen (Ref. 3710-001-K). HEK293T cells transfected to express murine NOG (HEK-NOG) were kindly provided by Dr. Muncan [21]. For each batch of CM, a cryovial of producer cells was thawed and expanded through passaging to obtain sufficient cell numbers. Cells were cultured in Dulbecco’s Modified Eagle Medium (DMEM) supplemented with GlutaMAX (Gibco, Ref 61965-026) and 10% heat-inactivated fetal bovine serum (HI-FBS) (Eurobio, Ref CVFSVF00-01). Cells were maintained at 37 °C in a humidified atmosphere with 5% CO_2_, medium was replaced every other day, and cells were passaged once they reached 80 to 90% confluence using a standard trypsin-EDTA protocol. During the expansion phase, selection antibiotics were applied for L-WNT3A, HEK-RSPO1, and HEK-NOG. This consisted of Geneticin (G418) (Invivogen, Ref. 108321-42-2; 400µg/mL), Zeocin (Invivogen, Ref. 11006-33-0; 300µg/mL) and Puromycin (Invivogen, Ref. 58-58-2, 10µg/mL), respectively.

For CM production, cells were harvested at near-confluence and counted using trypan blue exclusion to ensure accurate seeding densities: 0.7× 10□cells/mL for L WNT3A, 0.9 × 10 cells/mL for HEK-RSPO1, and 1 × 10 cells/mL for HEK-NOG. These defined seeding densities allowed cultures to reach confluence within 72 hours, corresponding to the first day of CM production. Upon reaching confluence, cultures were washed twice with serum-free DMEM + GlutaMAX to remove residual selection antibiotics. Cells were then incubated in fresh complete medium (DMEM + GlutaMAX + 10% HI-FBS) and CM was collected over two consecutive days following medium replacement, clarified by centrifugation (900 × g, 30 min, 4°C) to remove cellular debris, pooled, and aliquoted. Aliquots were subsequently frozen at −20°C and transferred to −80°C for long-term storage (up to 6 months).

### Conditioned media characterization

Characterization of the WNT3A-, RSPO1-, and NOG-CM was performed using different assays. *WNT3A*, *RSPO1, NOG* expression in the different cell lines were assessed using the procedures described in the RNA extraction and gene expression analysis section below. WNT3A concentration was quantified in the CM by ELISA using the Mouse Wnt-3a DuoSet ELISA kit (R&D Systems, Ref. DY1324B-05) according to the manufacturer’s instructions. For each assay, the basal CM production medium (DMEM + 1% GlutaMAX + 10% HI-FBS) was used as a negative control.

The functional activity of WNT3A-CM and RSPO1-CM was assessed using HEK-SuperTOPFlash (STF) reporter cells, which stably express luciferase under the control of a STF promoter containing TCF/LEF-responsive elements (kindly provided by Pr. Benoît Vanhollebeke, Laboratory of Neurovascular Signaling, ULB Neuroscience Institut, Belgium). HEK-STF cells were seeded in 96-well plates at a density of 3 × 10 cells/mL (100 µL per well) in DMEM/F-12 supplemented with GlutaMAX (Gibco, Ref 31331-028) and 10% HI-FBS. Cells were incubated at 37°C in a humidified atmosphere containing 5% CO-until reaching approximately 80% confluence. The following day, the medium was replaced with DMEM/F-12 + GlutaMAX supplemented with 2% HI-FBS to reduce basal signaling. After 24 h, cells were treated with WNT3A-CM alone, WNT3A-CM plus RSPO1-CM, or control medium, and incubated for an additional 24 h at 37°C and 5% CO_2_. To control potential differences in cell numbers, cell viability was assessed using the CellTiter-Fluor^TM^ Cell Viability Assay (Promega, Ref G6080) according to the manufacturer’s instructions. Fluorescence was measured using a microplate reader (TECAN). For luciferase activity measurement, the ONE-Glo^TM^ Luciferase Assay System reagent (Promega, Ref E6110) was used according to the manufacturer’s instructions, in the same culture wells. After the addition of the reagents, plates were incubated for several minutes at 37°C in the dark, and luminescence was subsequently measured using the microplate reader.

### Composition of in-house culture media

Two in-house culture media were used: the human intestinal organoid expansion medium (hIEM), used to maximize the generation of stem cell- and transit-amplifying cell-rich HIOs in Matrigel, in order to expand the lines and for biobanking purposes; and the human intestinal organoid differentiation medium (hIDM), used for HIOs differentiation. hIEM and hIDM were adapted from De Vries *et al.* [22], and Fujii *et al.* [23] respectively. Specifically, for hIDM, IGF-1 and FGF-2 were added to our medium to ensure the maintenance of complete cellular diversity directly after single-cell passage. hIEM and hIDM formulations, including actual volumes to obtain a ≈10 mL working aliquot, are reported in Table 1. A key resources list for the in-house media production is reported in Supplementary table 2.

**Table 1.**
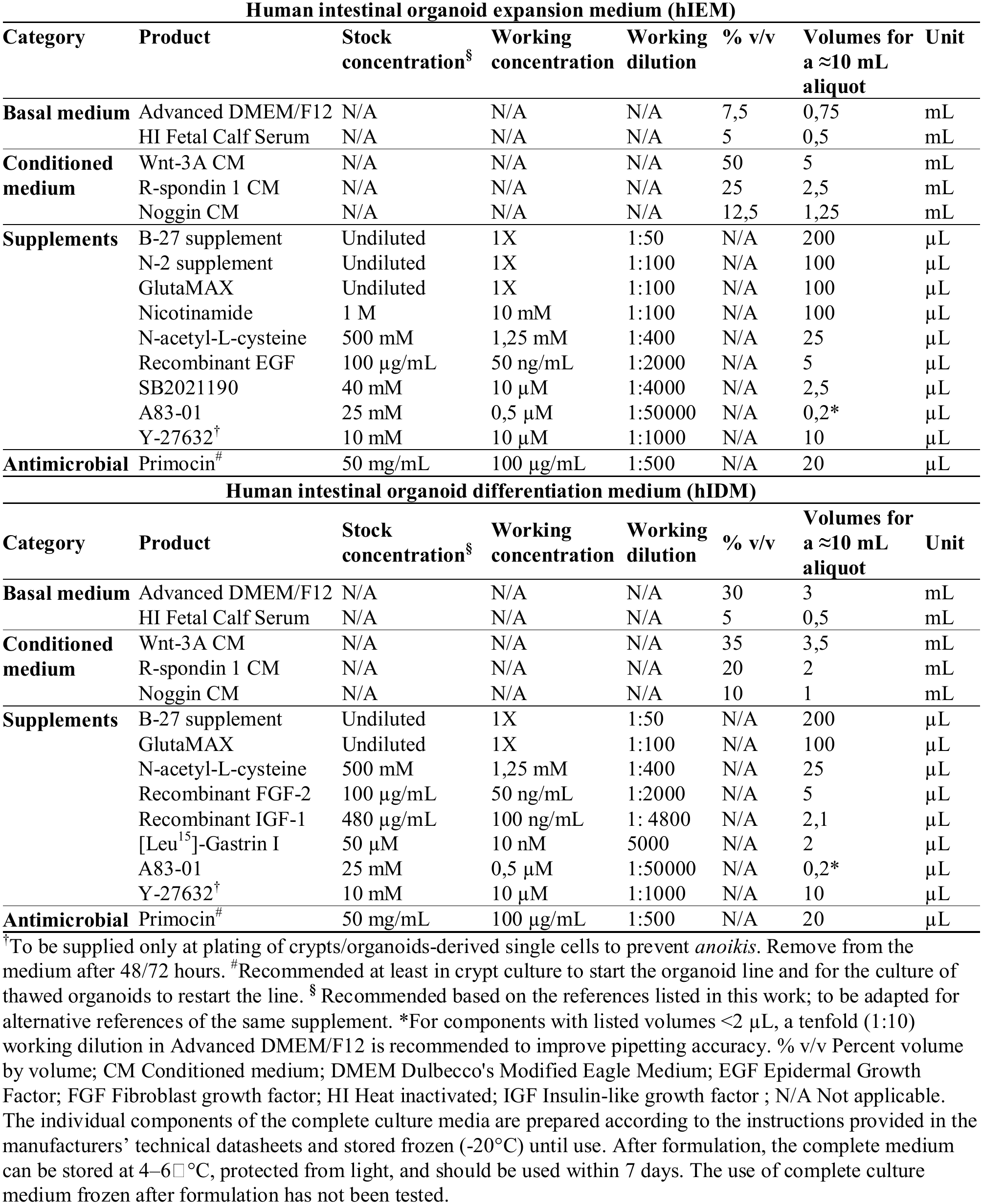
Organoid culture media formulations, including volumes for a single working aliquot.

| Human intestinal organoid expansion medium (hIEM) |  |  |  |  |  |  |  |
|---|---|---|---|---|---|---|---|
| **Category** | **Product** | **Stock concentration<sup>§</sup>** | **Working concentration** | **Working dilution** | **% v/v** | **Volumes for a ≈10 mL aliquot** | **Unit** |
| **Basal medium** | Advanced DMEM/F12 | N/A | N/A | N/A | 7,5 | 0,75 | mL |
|  | HI Fetal Calf Serum | N/A | N/A | N/A | 5 | 0,5 | mL |
| **Conditioned medium** | Wnt-3A CM | N/A | N/A | N/A | 50 | 5 | mL |
|  | R-spondin 1 CM | N/A | N/A | N/A | 25 | 2,5 | mL |
|  | Noggin CM | N/A | N/A | N/A | 12,5 | 1,25 | mL |
| **Supplements** | B-27 supplement | Undiluted | 1X | 1:50 | N/A | 200 | µL |
|  | N-2 supplement | Undiluted | 1X | 1:100 | N/A | 100 | µL |
|  | GlutaMAX | Undiluted | 1X | 1:100 | N/A | 100 | µL |
|  | Nicotinamide | 1 M | 10 mM | 1:100 | N/A | 100 | µL |
|  | N-acetyl-L-cysteine | 500 mM | 1,25 mM | 1:400 | N/A | 25 | µL |
|  | Recombinant EGF | 100 µg/mL | 50 ng/mL | 1:2000 | N/A | 5 | µL |
|  | SB2021190 | 40 mM | 10 µM | 1:4000 | N/A | 2,5 | µL |
|  | A83-01 | 25 mM | 0,5 µM | 1:50000 | N/A | 0,2* | µL |
|  | Y-27632<sup>†</sup> | 10 mM | 10 µM | 1:1000 | N/A | 10 | µL |
| **Antimicrobial** | Primocin<sup>#</sup> | 50 mg/mL | 100 µg/mL | 1:500 | N/A | 20 | µL |
| **Human intestinal organoid differentiation medium (hIDM)** |  |  |  |  |  |  |  |
| **Category** | **Product** | **Stock concentration<sup>§</sup>** | **Working concentration** | **Working dilution** | **% v/v** | **Volumes for a ≈10 mL aliquot** | **Unit** |
| **Basal medium** | Advanced DMEM/F12 | N/A | N/A | N/A | 30 | 3 | mL |
|  | HI Fetal Calf Serum | N/A | N/A | N/A | 5 | 0,5 | mL |
| **Conditioned medium** | Wnt-3A CM | N/A | N/A | N/A | 35 | 3,5 | mL |
|  | R-spondin 1 CM | N/A | N/A | N/A | 20 | 2 | mL |
|  | Noggin CM | N/A | N/A | N/A | 10 | 1 | mL |
| **Supplements** | B-27 supplement | Undiluted | 1X | 1:50 | N/A | 200 | µL |
|  | GlutaMAX | Undiluted | 1X | 1:100 | N/A | 100 | µL |
|  | N-acetyl-L-cysteine | 500 mM | 1,25 mM | 1:400 | N/A | 25 | µL |
|  | Recombinant FGF-2 | 100 µg/mL | 50 ng/mL | 1:2000 | N/A | 5 | µL |
|  | Recombinant IGF-1 | 480 µg/mL | 100 ng/mL | 1: 4800 | N/A | 2,1 | µL |
|  | [Leu<sup>15</sup>]-Gastrin I | 50 µM | 10 nM | 5000 | N/A | 2 | µL |
|  | A83-01 | 25 mM | 0,5 µM | 1:50000 | N/A | 0,2* | µL |
|  | Y-27632<sup>†</sup> | 10 mM | 10 µM | 1:1000 | N/A | 10 | µL |
| **Antimicrobial** | Primocin<sup>#</sup> | 50 mg/mL | 100 µg/mL | 1:500 | N/A | 20 | µL |

<sup>†</sup>To be supplied only at plating of crypts/organoids-derived single cells to prevent *anoikis*. Remove from the medium after 48/72 hours. <sup>#</sup>Recommended at least in crypt culture to start the organoid line and for the culture of thawed organoids to restart the line. <sup>§</sup> Recommended based on the references listed in this work; to be adapted for alternative references of the same supplement. \*For components with listed volumes <2 µL, a tenfold (1:10) working dilution in Advanced DMEM/F12 is recommended to improve pipetting accuracy. % v/v Percent volume by volume; CM Conditioned medium; DMEM Dulbecco's Modified Eagle Medium; EGF Epidermal Growth Factor; FGF Fibroblast growth factor; HI Heat inactivated; IGF Insulin-like growth factor ; N/A Not applicable. The individual components of the complete culture media are prepared according to the instructions provided in the manufacturers' technical datasheets and stored frozen (-20°C) until use. After formulation, the complete medium can be stored at 4–6□°C, protected from light, and should be used within 7 days. The use of complete culture medium frozen after formulation has not been tested.

During primary culture from isolated crypts and the first passage, IntestiCult™ Organoid Growth Medium Human (OGM) (STEMCELL Technologies, Ref. 06010) was used to optimize viability of the stem cells. Passages for HIOs expansion, biobanking, and characterization were performed in the in-house medium. All HIOs culture media used — hIEM, hIDM and OGM — were supplemented with Primocin (Invivogen, Ref ant-pm-1) to avoid the development of bacterial contaminations. Absence of mycoplasma contamination was tested regularly in cultures through third-party services (Eurofins, Ref 50400400).

### Crypt isolation and establishment of primary organoid cultures

Steps were adapted from Sato *et al.* [2]. Biopsies were transferred to 50 mL conical tubes and rinsed in cold DPBS. All subsequent steps were performed using pre-wet serological pipettes with an anti-adherence solution consisting of DPBS + 10% HI-SVF. To remove mucus and debris, tissue was briefly exposed to 10 mM dithiothreitol (Sigma-Aldrich, Ref 43816), immediately decanted, and washed in cold DPBS until the supernatant was clear. Epithelial crypts were isolated by incubating biopsies in 10 mM ethylenediaminetetraacetic acid (EDTA) (Invitrogen, Ref 15575020) in DPBS for 60 min on ice with gentle rocking (∼60 cycles/min). Mechanical dissociation was then performed by vigorous pipetting (≈10 passages through a 10 mL serological pipette) in cold DPBS, and the supernatant containing released crypts was collected. This dissociation step was repeated 2–5 times until adequate crypt yield was obtained. Crypt-containing fractions were centrifuged at 100 × g, 4 °C, 3 min to pellet crypts and pooled by being gently resuspended in ice-cold Matrigel (Corning, Ref CLS356231), plated in a 24-well tissue-culture treated plate (Sarstedt Ref 83.3922), by plating three 15 µL domes per well, and allowed to polymerize (20 min, 37°C, 5% CO2) before being overlaid with hIEM or OGM supplemented with Primocin (100 µg/mL) and Y-27632 (10 µM, STEMCELL Technologies, Ref #72304), the latter withdrawn after the first medium change. Cultures were maintained at 37°C, 5% CO2, with medium renewed every 48–72 h.

### HIOs expansion and differentiation

Following establishment of the primary HIOs cultures, lines were passaged on culture day ∼10 (range 7–12) for three generations when maintained in expansion condition, in order to preserve stemness. Culture medium was refreshed 24 hours prior to passaging to optimize viability. At each passage, approximately one-third of the HIOs were dedicated to continue the expansion of the line, while the remaining two-thirds were cryopreserved.

HIOs were recovered from Matrigel and washed several times using ice-cold Advanced DMEM/F-12 (Gibco, Ref 12634010), centrifugated (300 × g, 4°C, 5 min) and resuspended in 1 mL TrypLE Express (Gibco, Ref 12604013). Single-cell dissociation was achieved after a three steps incubation at 37°C (3 min, then 2 min, then 1 min), each time followed by mechanical trituration (10 passages through a nested P1000–P200 pipette-tip assembly). Enzymatic activity was then quenched by dilution in 10 mL of ice-cold Advanced DMEM/F-12, and single cells were pelleted by centrifugation (300 × g, 4°C, 10 min). To generate HIOs with differentiated epithelium, hIDM was applied from the first day of culture to single cells derived from the dissociation of HIOs previously expanded in hIEM. Viable single cells obtained after TrypLE dissociation were counted and their viability assessed by Trypan Blue exclusion using a Countess 3 FL automated counter. Cell pellets were resuspended in Matrigel at target densities depending on downstream application: ∼1000 cells/µL for continuing the expansion of the line, and ∼2000 viable cells/µL for epithelial differentiation for terminal use in downstream experiments. The higher seeding density for differentiation was intended to compensate for the expected cell loss under this condition and maintain adequate culture density for downstream experiments. Cell/Matrigel suspensions were plated as described above, overlaid with 450 µL of hIEM or hIDM – depending on the purpose of the culture – and supplemented at seeding with Y-27632 (10 µM), withdrawn after the first medium change.

### HIOs cryopreservation and biobanking

For biobanking, HIOs lines were cryopreserved applying predefined quality criteria: (i) good HIOs density in expansion phase (by visual inspection of the culture in transmitted light), (ii) early passage (≤ 3), and (iii) culture within day 7. Culture medium was refreshed 24 h before freezing to optimize viability, with 1–3 high-density wells typically yielding one cryovial. Following organoid recovery from Matrigel, pellets were gently resuspended in CryoStor® CS10 (STEMCELL Technologies, Ref #07931). Samples were kept on ice and controlled-rate freezing (−1°C/min) was initiated within 10 min of CS10 addition using a passive freezing container (Corning® CoolCell). Cryovials were held at −80 °C for 24 h before being transferred to liquid nitrogen for long-term storage.

### RNA extraction, and gene expression analysis by TaqMan assays

Total RNA was isolated from biopsies, isolated intestinal crypts, and HIOs using the NucleoSpin® RNA Mini kit (Macherey-Nagel, Ref 740955.250) according to the manufacturer’s instructions, including an on-column DNase step. RNA concentration was measured with a Qubit 3 Fluorometer and Qubit RNA Broad Range kit (Thermo Fisher Scientific, Ref Q10211). RNA integrity was assessed on a 2100 Bioanalyzer (Agilent Technologies). Typical yields/ RNA integrity score (RIN) was for biopsies 50–150 ng/µL, RIN > 7; for crypts 30–80 ng/µL, RIN > 8; for HIOs (one well) 100–250 ng/µL, RIN > 9. Complementary DNA (cDNA) was synthesized using the High-Capacity cDNA Reverse Transcription Kit (Applied Biosystems, Ref 4368814) according to the manufacturer’s instructions. cDNA was diluted 10 times and used immediately or transferred at −20 °C for deferred use. Gene expression was quantified on a StepOnePlus system (Applied Biosystems) using TaqMan™ FAM Gene Expression Assays (NFQ-MGB probes) and Takyon ROX Probe 2× MasterMix dTTP Blue (Eurogentec, Ref UF-RPMT-B0701). Each 20 µL reaction contained 10 µL MasterMix, 1 µL TaqMan assay, 1 µL cDNA, and 8 µL PCR-grade H_2_O, with reagents kept cold and protected from light, and plates briefly centrifuged before cycling. A detailed description of the TaqMan assays, comprising 2 housekeeping genes and 7 functional targets, along with the biological role of each marker, is provided in Supplementary table 3.

### Fluorescent immunostaining and imaging of HIOs and crypts

For microscopy studies, both thin histological sections from formaldehyde-fixed paraffin-embedded specimens and whole-mount samples for confocal and 3D light-sheet imaging were used. For immunofluorescent labeling of the histological preparations, samples were fixed in 4% weight-to-volume solution of paraformaldehyde (PFA) in DPBS (1 h, room temperature (RT)), washed in DPBS, and embedded in warm 2% agarose to stabilize the pellet. Agarose blocks were left to cure and stored in 70% ethanol (4–8°C) prior to automated dehydration, paraffin embedding, and 5 µm microtomy. Sections were deparaffinized according to standard procedures, subjected to antigen retrieval with a pH 6 citrate buffer (Dako Ref S2369; 98°C, 40 min), blocked with serum-free protein block (Dako Ref X0909; 60 min, RT), and incubated overnight (4–8°C) with primary antibody diluted in background-reducing diluent (Dako, Ref S3022). After washing (DPBS/0.1% Tween-20), sections were incubated with fluorophore-conjugated secondary antibodies (3 h, RT) and optional counterstain (DAPI, 15 min, RT; Hoechst, 15 minutes, RT; Phalloidin, 1 h, RT), washed, and mounted in ProLong™ Gold Antifade (Invitrogen, Ref P36930); coverslips were cured for 24 h before imaging.

For whole-mount 3D preparations, the procedure was adapted from a previously established protocol for high-resolution imaging of 3D organoids [24]. Samples were fixed in 4% PFA (1 h, RT), rinsed in DPBS, and sequentially permeabilized/blocked in DPBS/0.1% Tween-20 (60 min, RT) and DPBS/0.1% Triton X-100/0.2% BSA (washing buffer (WB)), using wide-bore, pre-wetted tips to minimize sample damage. Primary antibody solution (prepared in WB) was added at a 1:1 volume ratio per well (200 µL, low-attachment 24-well plates) and incubated overnight (4–8°C, mild rocking), followed by five washes in WB. For each wash step, the 24-well plate was tilted at a 45° angle, and HIOs were left to settle at the bottom of the well for 30 seconds; supernatant was subsequently aspirated. Secondary antibodies (3 h, RT) and optional counterstains (DAPI, 15 min, RT; Hoechst, 15 minutes, RT; Phalloidin, 1 h, RT) were applied following the same technique and wash cycles. Cleared whole-mounts were imaged promptly or stored at 4–8°C, with progressive fluorescence decay expected after ∼2 weeks. For light-sheet microscopy, fluorescently labeled samples were resuspended in 4% low-melting-point agarose (Invitrogen, Ref. 16520050) at 37°C and mounted in glass capillaries (Zeiss).

Imaging was performed at complementary levels of resolution and sophistication. Routine culture monitoring used an EVOS™ M5000 system (transmitted light and color-fluorescence mode, 4×–20× objectives) to document HIOs growth and morphology. Confocal microscopy was performed on a ZEISS LSM-700 (Plan-Apochromat 20×/0.8 NA objective), to image both immunofluorescent-labeled histological sections and whole-mounted samples, the latter using Z-stacks and tiled acquisitions (ZEISS Zen Blue). For volumetric imaging of larger specimens (whole-mounted HIOs up to ∼1 mm), light-sheet microscopy was performed on a ZEISS Lightsheet 7 (5× and 20× objectives) with labeled samples mounted in glass capillaries and embedded in refractive-index-matched clearing medium (60% glycerol/2.5 M fructose; RI ≈ 1.4688). A detailed list of antibodies and other fluorescent markers used with relative dilutions is reported in Supplementary table 4.

### Data processing and statistics

For RT-qPCR experiments, ΔCt of each sample was computed against a housekeeping reference (GUSB and/or HPRT1, as specified in each figure), and a calibrator condition (organoid grown in hIEM) was used to obtain ΔΔCt; foldchange was defined as 2^−ΔΔCt for visualization.

Statistical analyses were performed using R (version 4.2.3) in RStudio (version 2026.07.1+139). Non-parametric tests were applied using the rstatix package. For comparisons between two groups, the Wilcoxon rank-test (Mann-Whitney U test) was used. For comparisons among more than two conditions, an overall difference was first assessed using a Kruskal-Wallis test. When significant, pairwise multiple comparisons were performed using Dunn’s post hoc test, with p-values adjusted using the Benjamini-Hochberg (BH) method to control the false discovery rate (FDR). The specific statistical tests used are indicated in the figure legends. A p-value (or adjusted p-value for multiple comparisons) < 0.05 was considered statistically significant.

## RESULTS

### Levels and functionality of factors supporting HIOs culture in conditioned media

We first assessed the expression of *WNT3A*, *RSPO1*, and *NOG* in each engineered cell line across independent culture batches (Figure 1A). Our data confirmed that each cell line specifically expressed the corresponding factor, with expression levels remaining stable across independent culture batches. WNT3A protein quantification in L-WNT3A-CM through ELISA, showed a mean concentration of 470 ± 42 ng/mL, corresponding to approximately 235 ng/mL in the final hIEM (Figure 1B), with consistent WNT3A concentrations across independent production batches.

**Figure 1.**
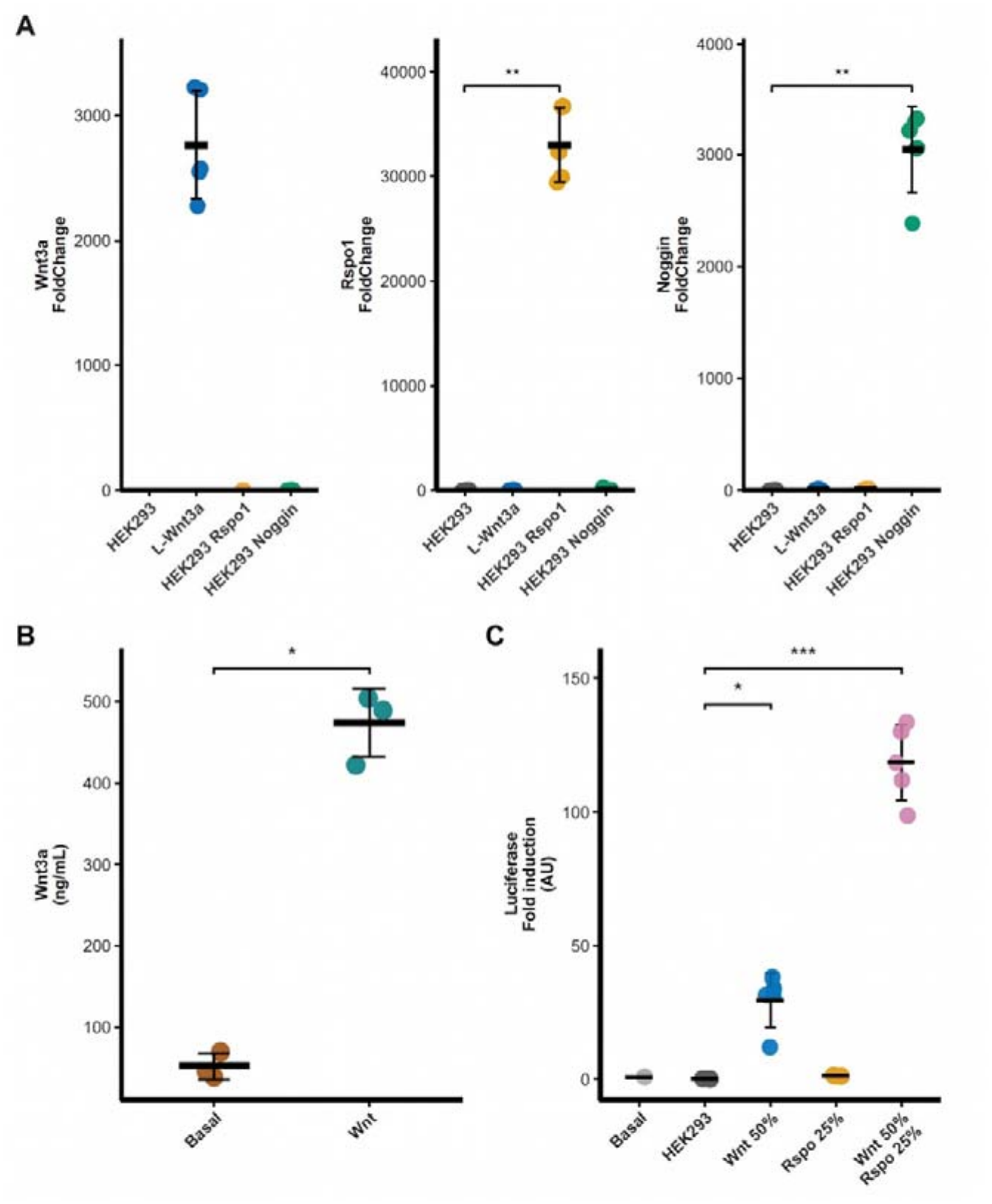
Characterization and functional analyses of in-house conditioned media for hIEM and hIDM formulation. **(A)** *WNT3A*, *RSPO1* and *NOG* expression levels in HEK293, L WNT3A, HEK-RSPO1 and HEK-NOG cell lines by RT-QPCR. Data is expressed as foldchange relative to HEK293 using the 2^−ΔΔCt^ method following normalization to *HPRT1* housekeeping gene; **(B)** Quantification of WNT3A in conditioned medium (CM) of L WNT3A cell line by ELISA assay. Basal: DMEM/HI-FBS; Wnt: WNT3A-CM; **(C)** Synergic activity of WNT3A/RSPO1-conditioned media (CM) assessed using the HEK293STF (SuperTOPFlash) luciferase reporter assay, following proportions used in hIEM medium. Cells were stimulated with WNT3A- and RSPO1-CM, and luciferase activity was normalized to cell viability measured with the CellTiter-Fluor assay. Basal: DMEM/HI-FBS; HEK293: HEK293-CM used as control; Wnt50%: WNT3A-CM diluted at 50%; Rspo25%: RSPO1-CM diluted at 25%. For panels **(A)** and **(C)**, statistical significance was assessed using the Kruskal– Wallis test followed by Dunn’s post hoc multiple-comparison test against the corresponding control in five independent batches. For panel **(B)**, statistical significance between the two groups was assessed using the Wilcoxon test in three independent batches. p < .05 (*), p < .01 (**), p < .001 (***).

For RSPO1, we assessed its biological activity in combination with WNT3A using a functional STF reporter assay. This assay evaluates the ability of the conditioned media to activate canonical Wnt/β-catenin signaling in HEK-STF reporter cells. Luciferase activity was normalized to cell viability measured using the CellTiter-Fluor assay to account for potential differences in cell number or metabolic activity. Treatment with WNT3A-CM alone induced a ≈30-fold increase in luciferase activity compared with basal culture medium. The addition of RSPO1-CM resulted in a further increase in reporter activity, up to ≈120-fold (Figure 1C). Finally, we confirmed the presence of Noggin protein in the CM by liquid chromatography coupled with mass spectrometry (Supplemental Table 5).

### The established culture conditions enable HIOs growth and expansion across donor groups

Starting from a standard caecal biopsy (Figure 2A), fresh crypts were isolated and plated in Matrigel (Figure 2B) to establish a new HIO line. The isolated crypts displayed preserved architecture and structural integrity, as assessed by whole-sample immunostaining for constitutive and functional markers (Figure 2C), including EpCAM as a pan epithelial marker, cytokeratin 20 and chromogranin A as markers of differentiated intestinal epithelium and Ki67 as a marker of cell proliferation. Crypts subsequently developed into three-dimensional budding structures (i.e. primary organoids) within the first days of culture (Figure 2D). On subsequent generation, HIOs maintained in hIEM exhibited a typical cystic morphology, characterized by a regular spherical shape and thin epithelial walls (Figure 2E). In contrast, HIOs cultured in hIDM promoted complex three-dimensional structure with a budding architecture, displaying contours reminiscent of the native intestinal epithelium and a darker, thickened lumen (Figure 2F). These structures reached several hundred micrometers in diameter within 10–12 days, with the largest HIOs approaching approximately 1 mm in diameter. HIO cultures were established and maintained similarly in all three donor groups (Supplementary figure 1). Taken together, this data indicates that HIOs cultures can be established and maintained in all three donor groups. Culture in hIEM enables repeated organoid passaging and long-term cryopreservation for biobanking and subsequent use.

**Figure 2.**
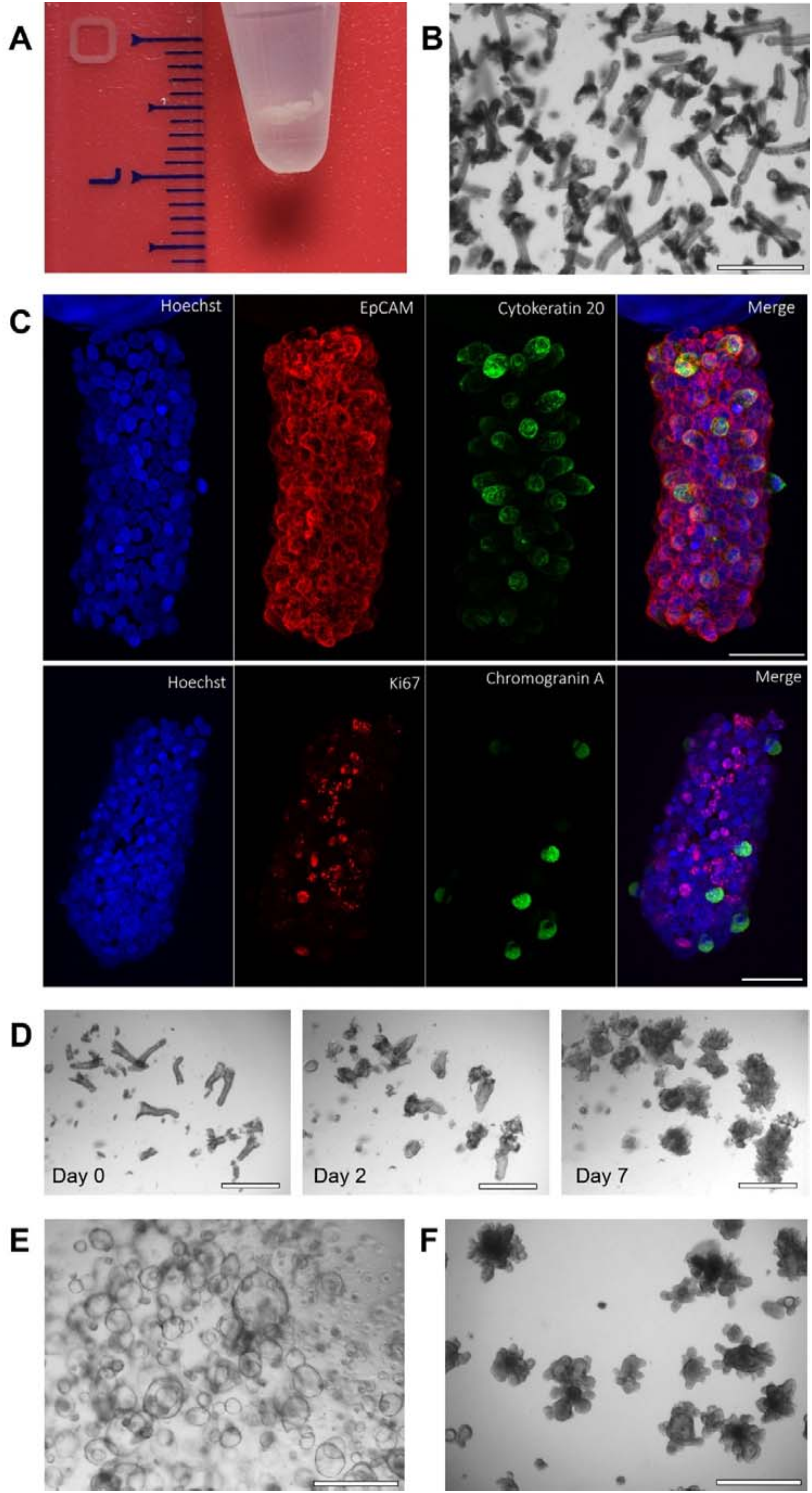
Sequential steps of the procedure from caecal biopsies to HIOs generation in hIEM and hIDM. **(A)** Human caecal biopsy (ruler in cm); **(B)** Caecal crypts freshly isolated from the biopsy; **(C)** Whole-mounted sample immunostaining of freshly isolated human caecal crypts: upper panels (left to right): Hoechst nuclear staining (blue), EpCAM (red), cytokeratin 20 (green), and merged acquisitions; lower panels (left to right): Hoechst nuclear staining (blue), Ki-67 (red), chromogranin A (green), and merged acquisitions; showed crypts are from the same donor (HC). **(D)** Morphological changes of crypts during the primary culture; budding enclosed structures are formed by day 7; showed primary culture is **(E)** Caecum HIOs grown in expansion medium (hIEM) and in **(F)** differentiation medium (hIDM); showed HIOs are from the same donor (SpA patient) and cultured in parallel in expansion vs differentiation condition (day 10 of culture). The acquisitions were performed on a ZEISS LSM-700 confocal microscope equipped with a Plan-Apochromat 20×/0.8 lens, scale bar = 50 μm **(C)** or in transmitted light using an EVOS M5000 microscope and a 4x objective **(B, D-F);** scale bar = 750 μm.

### Culture in hIDM rewires intestinal organoid programs toward a differentiated phenotype

To assess the effect of hIDM and compare the resulting phenotype with the initial tissue and expansion conditions, we examined the expression of *MUC2*, *KRT20*, and *CHGA* as markers of intestinal epithelial differentiation, together with *LGR5* (Figure 3) as a marker of intestinal stem cells. Gene expression was assessed across different sample types, including caecal biopsies, freshly isolated crypts, and HIOs cultured in either hIEM or hIDM.

**Figure 3.**
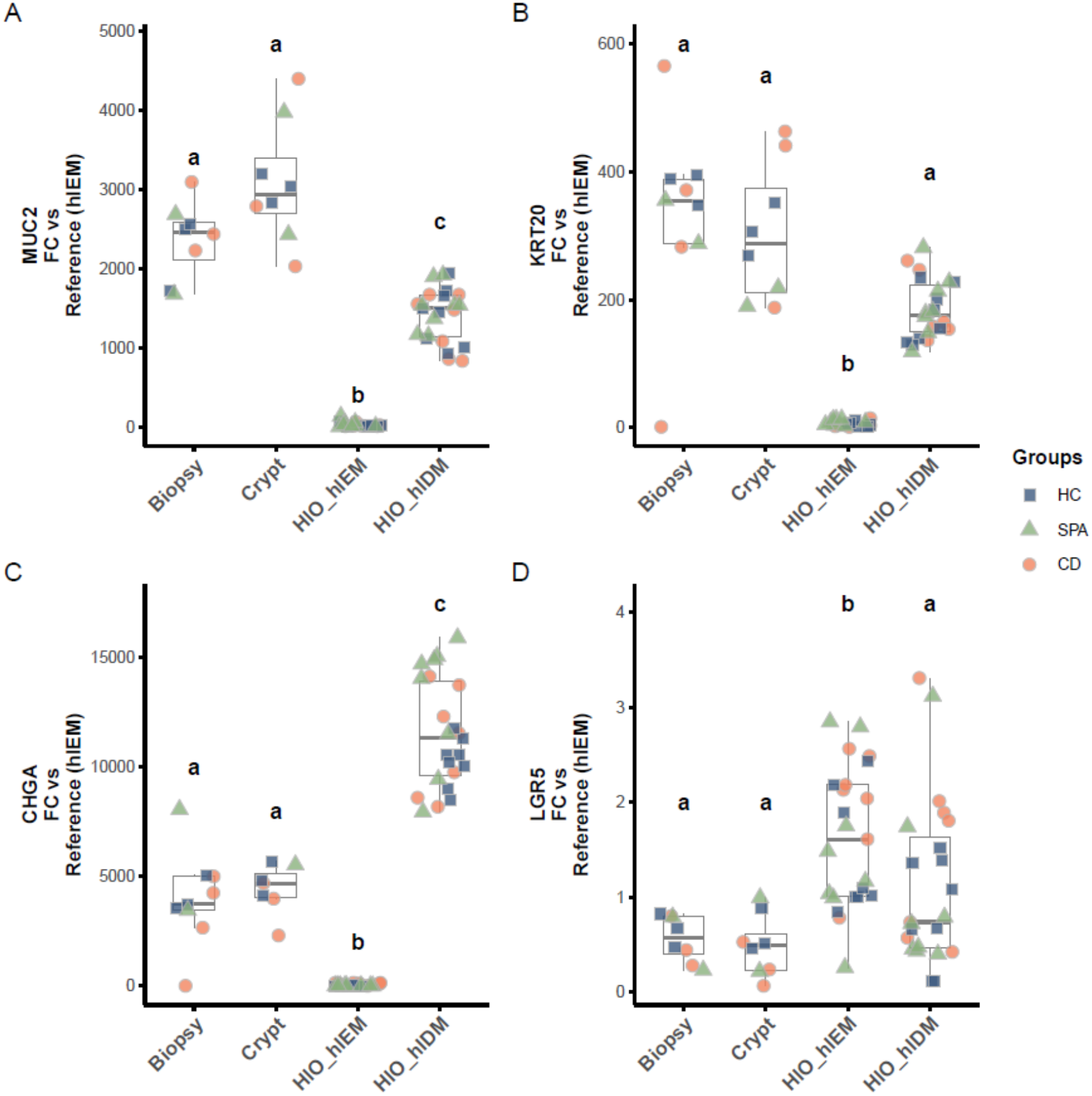
Expression of intestinal epithelial markers in caecal biopsies, isolated crypts, and HIOs grown in hIEM and hIDM. Caecal biopsies, isolated crypts, and HIOs grown in hIEM and hIDM were analyzed. Expression of **(A)** *MUC2* (mucin-2), **(B)** *KRT20* (cytokeratin 20), **(C)** *CHGA* (chromogranin A) and **(D)** *LGR5* was measured by RT-qPCR and expressed as foldchange (FC) relative to a hIEM sample (Reference) using the 2^−ΔΔCt^ method following normalization to *GUSB* housekeeping gene. Each point represents a sample, with colors and symbols indicating the donor group: Healthy Controls (blue square, HC), Spondyloarthritis (green triangle, SPA) and Crohn’s Disease (orange circle, CD). The data presented for biopsies, crypts, and HIOs were not always obtained from matched samples from the same donor, owing to limited tissue availability. Statistical significance was assessed using Kruskal-Wallis tests followed by Tukey’s post-hoc comparisons. Different letters (a, b, c) identify statistically significant differences between groups.

Gene expression analysis showed that in HIOs cultured in hIDM, *MUC2*, *CHGA*, and *KRT20* were upregulated showing – in contrast to what observed for HIOs cultured in hIEM - relative gene expression patterns skewed towards those of biopsies and crypts (Figure 3). These observations suggest that hIDM-cultured HIOs acquire a gene expression profile of pivotal epithelial markers resembling that of *in vivo* caecal epithelium, compared to an attenuated profile in standard expansion condition. Notably, *LGR5* expression was also detected in HIOs cultured in hIDM, although at lower levels than in HIOs maintained in hIEM. These results suggest that hIDM culture promotes the acquisition of differentiated epithelial features while maintaining detectable expression of the main intestinal stem cell compartment marker. Limited to the considered markers, the gene expression data were broadly overlapping across HIOs derived from HC-, SpA-, and CD-derived samples. Furthermore, the expression levels of *MUC2*, *KRT20*, *CHGA*, and *LGR5* were consistent across different culture batches, indicating stable gene expression profiles (Supplementary figure 2).

### Markers localization and proportion mimic that of tissue *in vivo*

Gene expression data were complemented by immunofluorescence staining performed either on histological sections or using 3D whole-mount imaging. Imaging analyses confirmed the presence and spatial distribution of multiple intestinal epithelial cell populations, including mucin 2 (MUC2)-positive goblet cells, chromogranin A (CHGA)-positive enteroendocrine cells, and cytokeratin 20 (CK20)-positive epithelial cells (Figure 4). CHGA-positive enteroendocrine cells were sparsely distributed throughout the HIOs, consistent with their relatively low abundance in the native intestinal epithelium (Supplementary figure 3). Ki-67-positive proliferative cells were predominantly localized at the base of the crypt-like structures (Figure 5A), particularly within the lower regions of the budding domains, consistent with the organization of the intestinal epithelium *in vivo* (Supplementary figure 3). MUC2-staining was predominantly oriented toward the luminal cavity (Figure 5B), consistent with the normal morpho functional organization of goblet cells.

**Figure 4.**
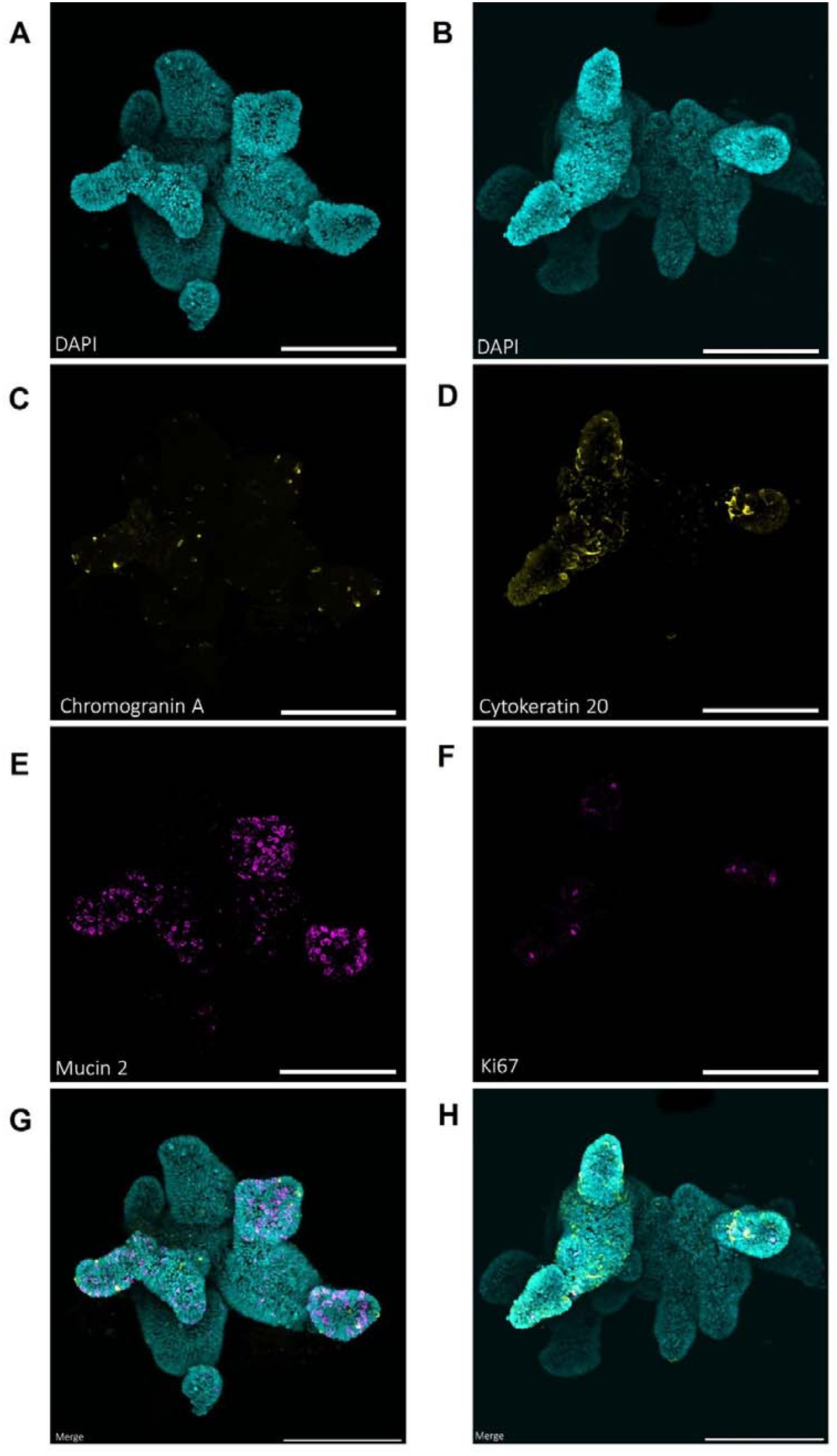
Differentiated HIOs display mature lineages while retaining proliferative potential. Immunostaining of hIDM-cultured HIOs reveals different intestinal epithelial lineages using whole-mount 3D imaging. **(A-B)** nuclear staining only (cyan); **(C)** Chromogranin A (yellow); **(D)** Cytokeratin 20 (yellow) **(E)** Mucin 2 (fuchsia); **(F)** Ki-67 (fuchsia); **(G-H)** merge of the above markers. Nuclei were counterstained with DAPI (blue). Figure shows SpA- (left panels) and HC-derived (right panels) caecum HIOs grown in hIDM and collected at day-10 of culture. The samples were fructose–glycerol-cleared and whole-mounted on a glass slide. The acquisition was performed on a ZEISS LSM-700 confocal microscope equipped with a Plan-Apochromat 20×/0.8 lens. Three-dimensional reconstruction was performed starting from a Z-stack and 2 x 2-tiled collection in Zeiss ZEN Blue. Scale bar = 200 µm.

**Figure 5.**
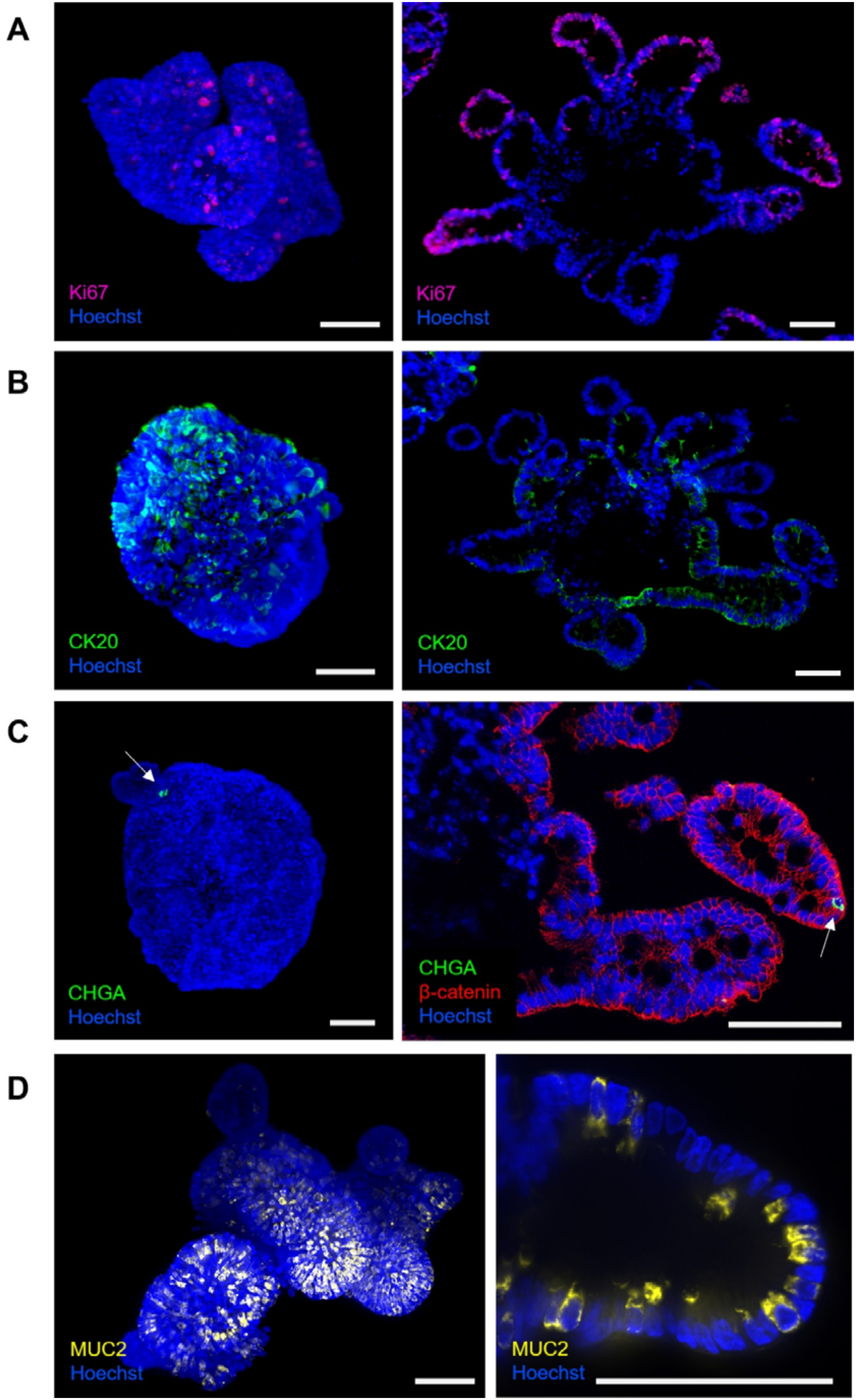
Spatial patterning of specific epithelial lineages in differentiated HIOs. Immunostaining of hIDM-cultured HIOs reveals the spatial distribution of multiple intestinal epithelial lineage markers. **(A)** Ki-67 (fuchsia); **(B)** Cytokeratin 20 (green); **(C)** Chromogranin A (green, highlighted by white arrows) and β-catenin (red; only tested for the sample shown in the right panel); **(D)** Mucin 2 (yellow). Nuclei were counterstained with Hoechst (blue). Figure shows HC- (left panels of A, B and C; both panels of D) and CD-derived (right panels of A, B and C) caecum HIOs grown in hIDM and collected at day-10 of culture. The acquisition of 3D whole-mount samples (left panels of A, B and C; both panels of D) was performed on a ZEISS Lightsheet 7 microscope equipped with a 5×/0.16 (left panels of A, B, C and D) or a 20×/1.0 detection lens (right panel of D), after fructose–glycerol-clearing, and volumetric reconstruction was carried out by Multiview mode (ZEN Blue). Histological sections (right panels of A, B and C) were imaged using an EVOS M5000 microscope in color-fluorescence mode and a 20× objective. Scale bar = 80 µm (all panels)

Structural and polarity-resolving markers, including F-actin, villin, E-cadherin and zonula occludens-1 (ZO-1), further highlighted the apical-in organization of the epithelium with strong preservation of epithelial apico-basal polarity (Supplementary figure 4).

Together, these findings demonstrate that hIDM-cultured HIOs develop a differentiated epithelial architecture comprising multiple intestinal cell lineages with distinct spatial distributions and preserved epithelial polarization, recapitulating key organizational features of the native intestinal epithelium.

## DISCUSSION

Here, we describe an integrated workflow to generate, expand, biobank, and differentiate HIOs from endoscopic biopsies of caecum, using two in-house media. The feasibility of our workflow was tested across a continuum of intestinal inflammation, clinically spanning from virtually absent inflammation in HCs, to subclinical/mild inflammation in SpA patients [19] and overt inflammation in CD patients [20], allowing the establishment of a clinically annotated biobank of human-derived epithelial lines. Our findings, based on the comparison of HIOs with biopsies and crypts fit within a growing body of evidence showing that, in selected preclinical contexts, HIOs can effectively recapitulate some valuable characteristics of the *in vivo* human epithelium [25–27]. Given the central role of the intestinal epithelium in tissue function and its frequent involvement in several diseases [28,29], recapitulating epithelial cellular diversity *in vitro* is essential for both fundamental mechanistic studies and translational applications.

To our knowledge, this is the first study to formally demonstrate the generation of HIOs from patients with SpA. The generation of SpA-derived HIOs extends this workflow to a donor group relevant to gut–joint axis research [20,30,31].Given the contribution of host genetics to SpA susceptibility [32,33], donor-derived epithelial cultures may support future mechanistic studies in a patient-specific genetic context. Their value for disease-focused and therapeutic research requires dedicated investigation [30,34]. These opportunities address an important unmet need in a chronic inflammatory disease associated with substantial functional impairment and reduced quality of life, for which no curative treatment is currently available.

In our study, HIOs lines were established exclusively from caecal tissue, allowing sampling to be standardized to a specific, well-defined anatomical zone. This approach limits a major source of inter-sample heterogeneity, namely the variability linked to position along the proximal-distal axis of the intestine, as the regional identity of the epithelium is known to be intrinsically programmed and stable in culture [35]. This has notably been confirmed in humans, where region-specific DNA methylation signatures are retained in intestinal organoids over extended culture periods [18].

Building on established organoid culture components [23,36], in which a combination of EGF, IGF-1, and FGF-2 enabled HIOs expansion while preserving cellular diversity, we described and tested two defined, in-house media - hIEM (expansion) and hIDM (differentiation) - for propagating and differentiating HIOs from HCs and diseased donors. The disclosed composition of in-house media allows individual components to be rationally titrated, and their contribution to specific epithelial readouts to be assessed, thereby favoring cross-laboratory reproducibility and tunability of the system [9,10].

We chose to work with CM for WNT3A, RSPO1, and NOG. Indeed, it has been shown that, at least for WNT3A, media supplemented with recombinant WNT3A alone failed to support the long-term survival of human colon organoids [11], in contrast to conditioned-medium formulations, underscoring the added value of CM over purified recombinant components. Furthermore, we chose to produce each CM individually rather than using the traditional L-WRN cell line [37], in order to be able to modulate WNT3A, RSPO1, and NOG independently. This approach also offers a more cost-effective source of these factors compared to purified recombinant proteins.

A recurrent limitation in the field is that CM are typically dosed by volume - most protocols use CM at a standardized dilution (commonly 50% for WNT3A-CM for example) - rather than by quantified protein concentration or functional activity. This issue was previously reported for the L-WRN cell line, for which exact WNT3A, RSPO1, and NOG concentrations could not be reliably determined by direct quantification techniques; batch potency was instead assessed functionally, through organoid growth assays or Wnt reporter assays [37].

To address this limitation, we implemented a workflow to characterize CM used by combining gene expression, ELISA, mass spectrometry and functional bioassays for our WNT3A-, RSPO1-, and NOG-CMs. ELISA showed that our WNT3A-CM contained 400-500 ng/mL of WNT3A. Gene expression analysis confirmed increased transcript levels of *WNT3A*, *RSPO1*, and *NOG* in the corresponding engineered cell lines. Functional activity was confirmed using the STF reporter assay, with magnitude of pathway activation comparable to that reported in previous studies using biologically active recombinant or conditioned WNT3A/RSPO1 preparations [36]. RSPO1-CM strongly potentiated WNT3A signaling. Furthermore, batch-to-batch activity remained reproducible, consistent with a multi-laboratory study in L-WRN-derived CM [38].

Across donors, HIOs cultured under our conditions consistently exhibited changes in growth, morphology, and gene expression indicative of epithelial maturation. These findings were supported by complementary analyses, including targeted intestinal gene RT-qPCR, and imaging in bright-field and after the immunostaining of lineage-specific antigens. Notably, differentiation condition induced a clear morphological transition from a predominantly cystic architecture to budding structures, resembling aspects of the native crypt–villus organization. This morphological remodeling was accompanied by changes in the expression and distribution of intestinal lineage markers, consistent with epithelial differentiation.

Our data confirm that HIOs mimic selected characteristics of the tissue they originate from. This capability has been reported across human organoid models: jejunal, ileal, and colonic organoids preserve the expression of genes specific to their segment of origin, as well as the core epithelial differentiation programs, over numerous passages in culture [39]. Similar findings have been obtained using single-cell transcriptomic approaches, which have shown that organoid-derived epithelial populations broadly recapitulate the cellular diversity observed *in vivo* [9,40].

Regardless of donor group, HIOs were successfully established from isolated crypts, with no difference in establishment success between patient- and healthy donor-derived samples. This contrasts with previous studies reporting lower establishment rates for IBD-derived organoids, particularly when biopsies were collected from inflamed intestinal regions [41]. Similarly, although altered organoid growth has been reported in IBD [42,43], we observed no significant difference in HIO growth rate between patient- and healthy donor-derived cultures. Importantly, in our study, biopsies were systematically collected from macroscopically unaffected intestinal regions. This sampling strategy may potentially explain the comparable organoid establishment and growth observed between patients and healthy donors in this work.

Formal comparisons between donor groups were beyond the scope of this study, which was not powered to detect intergroup differences. Nevertheless, proliferation and differentiation markers showed comparable expression levels and localization across all donor groups. Similar *MUC2*, *CHGA*, *KRT20*, and *LGR5* expression across groups was consistent with that observed in the original tissue (biopsies or crypts), with no apparent differences across the cohort.

## CONCLUSION

Taken together, our results confirm that HIOs cultured in hIDM recapitulate key molecular and morphological features of differentiated intestinal epithelium, irrespective of whether they are derived from healthy or diseased donor tissue.

Beyond modeling normal intestinal function, patient-derived organoids provide a valuable platform for investigating disease-associated epithelial phenotypes, when supported by appropriate experimental validation [7,42–49]. The rapidly growing number of studies reporting the successful establishment of large-scale, patient-derived living biobanks reflects the increasing recognition of their translational value. In this context, the inclusion of SpA -derived lines provides a novel resource for future studies of epithelial biology and host–microbiota interactions. Disease-specific mechanisms and the prediction of treatment responses remain questions for dedicated studies.

## Funding sources

This work was supported by the French Agence Nationale de la Recherche funded project MICROSPA and in the framework of the Carnot Qualiment© (doi.org/10.17180/h5gd-gk88) funded project Model Gut (https://qualiment.fr/des-projets-pour-anticiper-les-besoins-de-recherche-des-entreprises-agroalimentaires-2022/hot-topics/), and by Foundation for Research in Rheumatology (FOREUM) grant #069/2022 granted to G.N.

## Acknowledgements

The authors would like to thank Pierre Adenot and Matthieu Simion for their support on microscopy observations and access to the MIMA2 platform (Microscopie et Imagerie des Microorganismes, Animaux et Aliments, https://doi.org/10.15454/1.5572348210007727E12) as well as Marthe Vilotte and Julie Rivière for their help with histological preparations at the @BRIDGe histology facility https://abridge.inrae.fr) and for the analyzes carried out and the ISC CRB-Anim (https://doi.org/10.15454/1.5613785622827378E12). We would also like to thank Paloma Piquet and François Becher at the SPI/LI-MS facilities at the CEA in Paris Saclay for the liquid chromatography analyses. We sincerely thank all the patients who participated in this study for their time, trust, and valuable contribution to this research. Figures were generated using BioRender (https://biorender.com).

## Author contributions

GN : Writing – original draft, Conceptualization, Formal analysis, Funding acquisition; RS : Methodology, Conceptualization, Formal analysis; CLP : Methodology, Resources, Data curation; PD : Methodology, Data curation, Resources; AB : Resources; RSN : Methodology, Resources; MADA: Funding acquisition, Conceptualization; MB : Funding acquisition, Conceptualization, Supervision; TB : Funding acquisition, Conceptualization, Supervision; CC : Funding acquisition, Supervision, Conceptualization, Writing – review and editing. All authors have read and agreed to the published version of the manuscript.

## Declaration of competing interest

The authors declare no competing interest.

## Data availability

All data sets in this study are available from the corresponding authors upon reasonable request.

## Declaration of generative AI and AI-assisted technologies in the manuscript preparation process

The authors used OpenAI’s GPT-6 Astra and earlier GPT models for assistance with code development, language editing, phrasing refinement, and manuscript structure and presentation. All AI-assisted code, analytical outputs, and text were reviewed and validated by the authors. The authors retained responsibility for experimental design, analytical choices, literature evaluation, interpretation of results, and final scientific and editorial decisions, and take full responsibility for the accuracy, integrity, and originality of the manuscript, including its data, figures, and conclusions.

## Notes

### Competing Interest Statement

The authors have declared no competing interest.

