## Supplemental material for "Human intestinal organoids from spondyloarthritis, Crohn’s disease and healthy controls: an integrated workflow for expansion, biobanking, differentiation, and characterization"

### SUPPLEMENTARY MATERIAL

#### Supplementary table 1. Inclusion and exclusion criteria

| **Inclusion criteria** | |
| --- | --- |
| **Population** | **Definition** |
| General eligibility (all groups) | Adult participants with a clinical indication for colonoscopy, including colorectal cancer screening. |
|  | Capacity to understand the study information and provide informed consent, with voluntary written informed consent obtained before any study-specific procedure. |
| Spondyloarthritis group | Patients fulfilling the ASAS classification criteria for axial and/or peripheral SpA [49] |
| Healthy control group | Individuals without clinically apparent gastrointestinal or systemic disease on clinical assessment, undergoing colonoscopy as part of a screening programme. |
| Crohn’s disease group | Established diagnosis of Crohn’s disease, supported by documented endoscopic findings and compatible histopathology. |
| **Exclusion criteria** | |
| **Condition** | **Definition** |
| Malignancy | Known or suspected gastrointestinal malignancy, including colorectal cancer, or active extraintestinal malignancy under treatment. |
| Celiac disease | Established diagnosis of celiac disease. |
| Gastrointestinal anatomy or obstruction incompatible with standardized caecal sampling | Previous right hemicolectomy, ileocaecal resection, or stoma; or intestinal strictures preventing caecal intubation. |
| Inadequate bowel preparation | Bowel preparation insufficient to permit reliable mucosal assessment. |
| Infectious or severe colitis | Suspected or confirmed infectious colitis, including *Clostridioides difficile* infection; or severe colitis or toxic megacolon requiring hospitalization. |
| Recent systemic antibiotic exposure | Systemic antibiotic use within the 4 weeks preceding colonoscopy. |
| Severe uncontrolled comorbidity or procedural contraindication | Hemodynamic or respiratory instability, or any other condition rendering colonoscopy or sedation unsafe, as assessed by the endoscopist. |

#### Supplementary table 2. Key resources list for the in-house media.

| **Reagent or resource** | **Origin** | **Reference** |
| --- | --- | --- |
| **Culture media, buffers, general chemicals** |  |  |
| Advanced DMEM/F-12 | Gibco | 12634028 |
| Fetal calf serum (South America) | Eurobio Scientific | CVFSVF00-01 |
| GlutaMAX Supplement | Gibco | 35050061 |
| B-27 Supplement (50X), serum-free | Gibco | 17504044 |
| N-2 Supplement (100X) | GIbco | 17502048 |
| N-Acetyl-L-cysteine, cell-culture grade | Sigma-Aldrich | A9165 |
| Primocin, antimicrobial agent for primary cells | Invivogen | ant-pm-1 |
| **Peptides, recombinant proteins, small molecules** |  |  |
| [Leu^15^]-Gastrin I human | Sigma-Aldrich | G9145 |
| Animal-Free Recombinant Human EGF | Peprotech | AF-100-15 |
| Recombinant Human IGF-1 (carrier-free) | BioLegend | 590908 |
| Recombinant Human FGF-2 (basic) (154 a.a.) | Peprotech | 100-18B |
| Y-27632 | STEMCELL Technologies | 72304 |
| A83-01 | STEMCELL Technologies | 72024 |
| SB2021190 | STEMCELL Technologies | 72632 |

#### Supplementary table 3. List of TaqMan probes (FAM-MGB) used in the study.

| **Target symbol** | **Assay ID** | **Gene Name** | **Description** |
| --- | --- | --- | --- |
| **GUSB** | Hs00939627_m1 | Glucuronidase beta | Housekeeping |
| **HPRT1** | Hs02800695_m1 | Hypoxanthine phosphoribosyltransferase 1 | Housekeeping |
| **MUC2** | Hs03005094_m1 | Mucin 2 | Epithelial differentiation/  Goblet cell |
| **KRT20** | Hs00300643_m1 | Cytokeratin 20 | Epithelial differentiation/  Absorbitive cell |
| **CHGA** | Hs00900370_m1 | Chromogranin A | Epithelial differentiation/  Enteroendocrine cell |
| **LGR5** | Hs00969422_m1 | Leucine rich repeat containing G protein-coupled receptor 5 | Stem cell niche |
| **WNT3A** | Mm00437337_m1 | Wingless-type MMTV integration site family, member 3A | Stem cell / Progenitor proliferation, differentiation inhibition |
| **RSPO1** | Mm00507077_m1 | R-spondin 1 | Stem cell / Progenitor proliferation, differentiation inhibition |
| **NOG** | Mm01297833_s1 | Noggin | Stem cell / Progenitor proliferation, differentiation inhibition |

#### Supplementary table 4. List of antibodies used in the study for immunofluorescence experiments

| **Antigen** | **Clone** | **Conjugate** | **Species** | **Supplier** | **Reference** | **Dilution for WMS** | **Dilution for HP** |
| --- | --- | --- | --- | --- | --- | --- | --- |
| **Primary ABs** | | | | | | | |
| Ki-67 | MIB-5 | Unconjugated | Mouse | Dako | M7248 | 1:100 | 1:200 |
| Mucin 2 | CCp58 | Unconjugated | Mouse | Santa Cruz | sc-7314 | 1:200 | 1:300 |
| Β-catenin | E-5 | Unconjugated | Mouse | Santa Cruz | sc-7963 | 1:200 | 1:300 |
| Chromogranin A | Polyclonal | Unconjugated | Rabbit | Abcam | ab45179 | 1:200 | 1:300 |
| Cytokeratin 20 | Sp33 | Unconjugated | Rabbit | Abcam | ab64090 | 1:200 | 1:300 |
| EpCAM | Polyclonal | Unconjugated | Goat | Novus | AF960-SP | 1:300 | 1:300 |
| Zonula occludens 1 | 1A12 | Unconjugated | Mouse | Invitrogen | 33-9100 | 1:200 | 1:200 |
| Villin | SP145 | Unconjugated | Rabbit | Abcam | ab130751 | 1:200 | 1:300 |
| E-cadherin | Polyclonal | Unconjugated | Goat | Novus | AF748-SP | 1:300 | 1:300 |
| **Secondary ABs** | | | | | | | |
| Rabbit IgG | Polyclonal | Alexa Fluor 488 | Donkey | Invitrogen | A-21206 | 1:2000 | 1:2000 |
| Goat IgG | Polyclonal | Alexa Fluor 647 | Donkey | Invitrogen | A-21447 | 1:2000 | 1:2000 |
| Mouse IgG | Polyclonal | Alexa Fluor 555 | Donkey | Invitrogen | A-31570 | 1:2000 | 1:2000 |
| **Other reagents** | | | | | | | |
| Hoechst 33342 | N/A | N/A | N/A | Invitrogen | H3570 | 1:2000 | 1:2000 |
| DAPI | N/A | N/A | N/A | Invitrogen | D3571 | 1:1000 | 1:1000 |
| F-actin (phalloidin) | N/A | iFluor 647 | N/A | Abcam | ab176759 | 1:1000 | 1:1000 |
| WMS Whole-mounted sample; HP Histological preparation; N/A Not applicable | | | | | | | |

#### Supplementary table 5. Mass spectrometric identification of Noggin in conditioned medium.

| **Peptide sequence** | **Reported modifications** | **PSMs: replicate 1** | **PSMs: replicate 2** | **Reported confidence** |
| --- | --- | --- | --- | --- |
| CGWIPIQYPIISECK | 2 × carbamidomethylation | 4 | 5 | High |
| GLEFSEGLAQGK | None reported | 3 | 3 | High |
| LQMWLWSQTFCPVLYAWNDLGSR | 1 × carbamidomethylation | 3 | 2 | High |
| QRPSGAMPSEIK | None reported | 3 | 3 | High |
| SCSVPEGMVCKPSK | 2 × carbamidomethylation | 1 | 1 | High |
| SLLGGHYDPGFMATSPPEDRPGGGGGPAGGAEDLAELDQLLR | None reported | 3 | 2 | High |
| SVHLTVLR | None reported | 2 | 3 | High |

Noggin-conditioned medium in serum-free DMEM was analyzed in two replicates (Rep1 and Rep2). Both identified the same seven unique peptides assigned to Noggin (UniProt P97466, *Mus musculus*), covering 54% of its sequence, with 19 peptide–spectrum matches (PSMs) per replicate. Sequest scores were 77.69 and 77.58 for replicates 1 (Rep1) and 2 (Rep2), respectively.

Liquid chromatography coupled with electrospray ionization tandem mass spectrometry (LC-ESI-MS/MS) was performed using a Q Exactive Hybrid Quadrupole-Orbitrap mass spectrometer operated in positive ionization mode with a data-dependent acquisition method selecting the top 10 precursor ions for fragmentation. Raw data were analyzed using Proteome Discoverer v2.5, with database searches performed against either the mammalian proteome sequences available in UniProt or a predefined list of selected proteins. The search parameters included trypsin as the proteolytic enzyme, N-terminal acetylation and methionine oxidation as variable modifications, and cysteine carbamidomethylation as a fixed modification. The precursor and fragment mass tolerances were set to 10 ppm and 0.02 Da, respectively, and up to two missed cleavages were allowed.

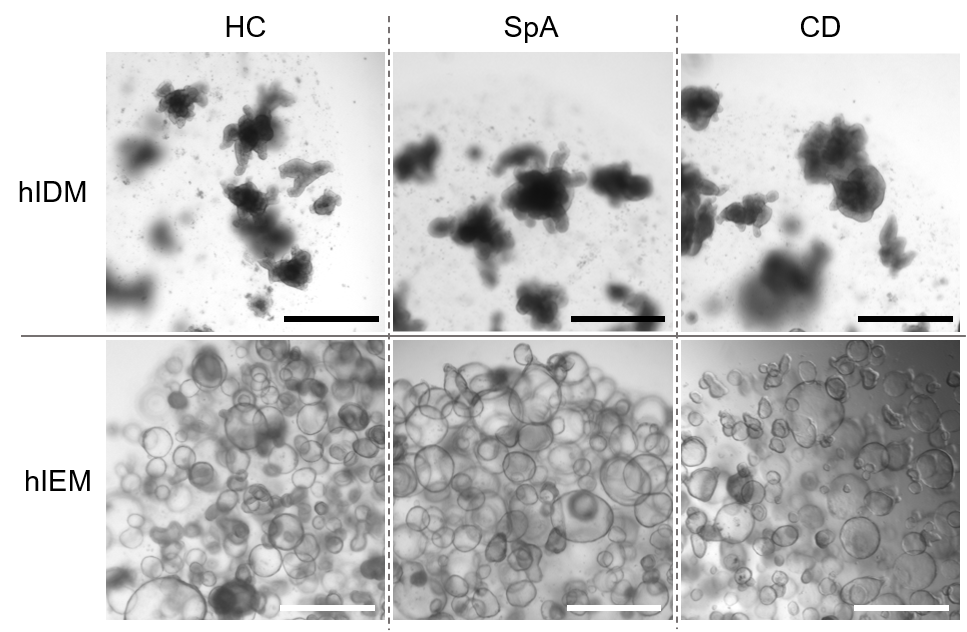

#### Supplementary figure 1. General morphology of caecal HIOs derived from different subgroups.

Representative images showing no major differences in general morphology of HIOs derived from donors from the three subgroups both in expansion and differentiation conditions. Each column shows HIOs from the same donor cultured in parallel in expansion vs differentiation condition (day 10 of culture). The acquisitions were performed in transmitted light using an EVOS M5000 microscope and a 4x objective. Scale bars = 750 μm.

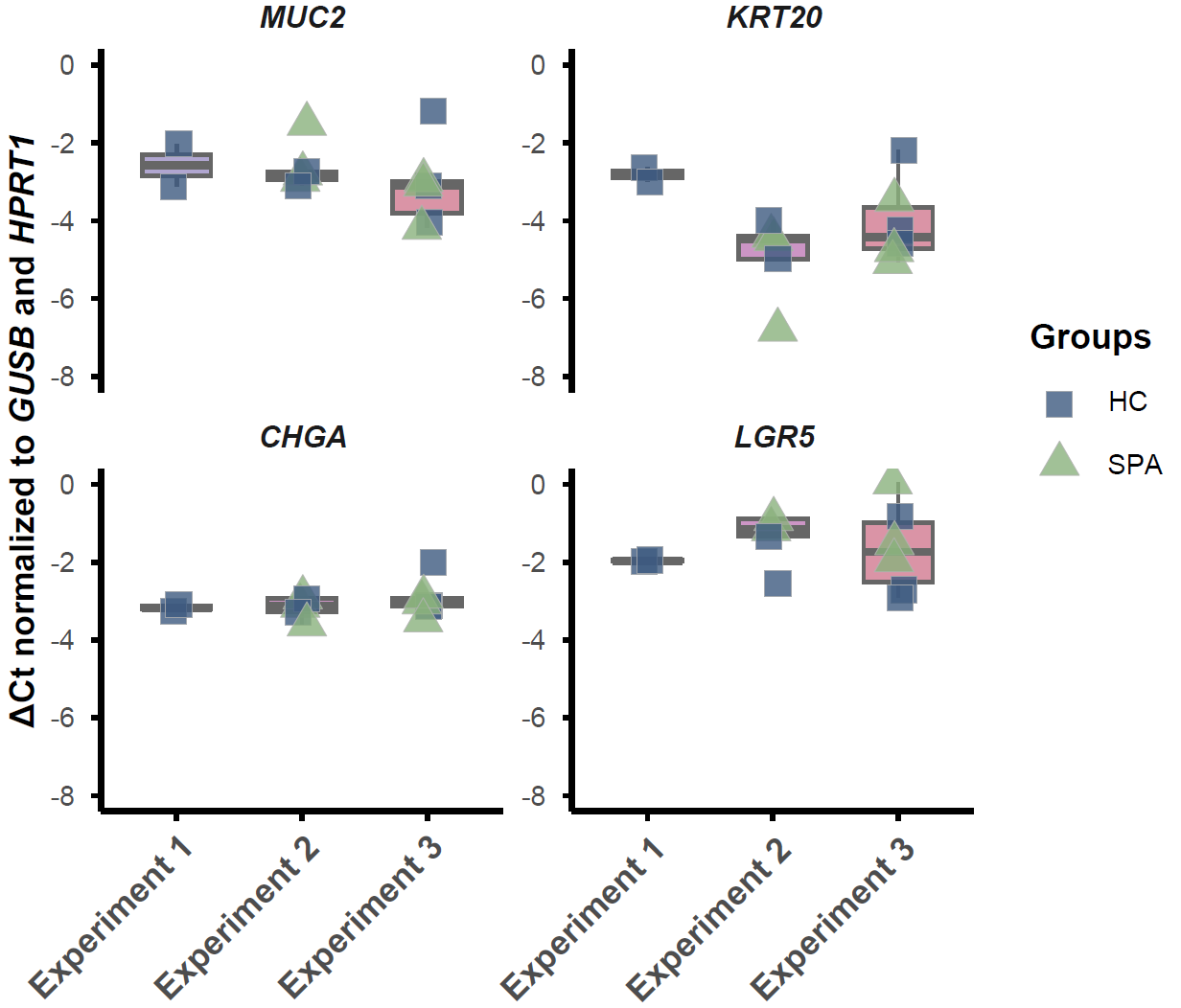

#### Supplementary figure 2. Expression of intestinal epithelial markers in HIOs grown in hIDM across different experimental runs.

HIOs grown in different batches of hIDM were analyzed in three separate experimental runs. Expression of MUC2 (mucin-2), KRT20 (cytokeratin 20), CHGA (chromogranin A), LGR5 was measured by RT-qPCR and expressed as ΔCt normalized to GUSB and HPRT1 housekeeping gene. Each point represents a sample, with colors and symbols indicating the donor group: Healthy Controls (blue square, HC) and Spondyloarthritis (green triangle, SPA).

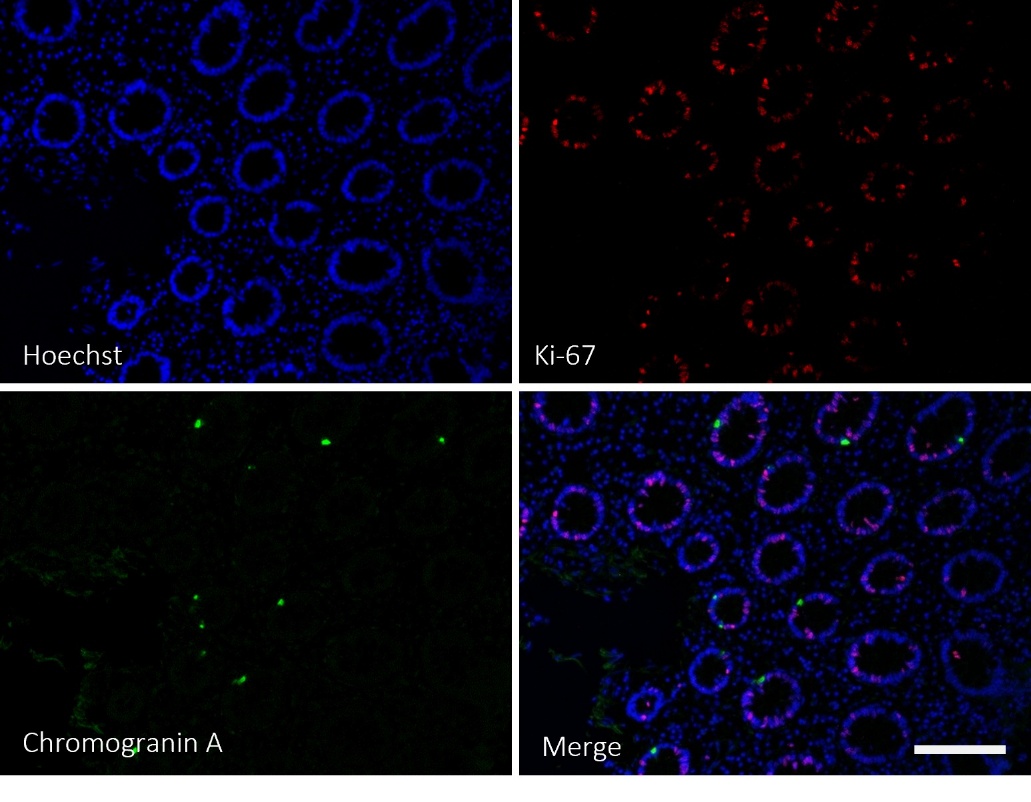

#### Supplementary figure 3. Immunofluorescence staining of native human epithelium.

Immunostaining of histological preparation of human caecal biopsies showing relative abundances and spatial distribution of intestinal functional markers Ki-67 (red, upper right panel) and chromogranin A (green, lower let panel). Nuclei were counterstained with Hoechst (blue) and merged with other acquisitions (lower right panel). Figure shows a SpA-derived sample. The acquisition was performed using an EVOS M5000 microscope in color-fluorescence mode and a 20× objective. Scale bar = 80 µm (all panels).

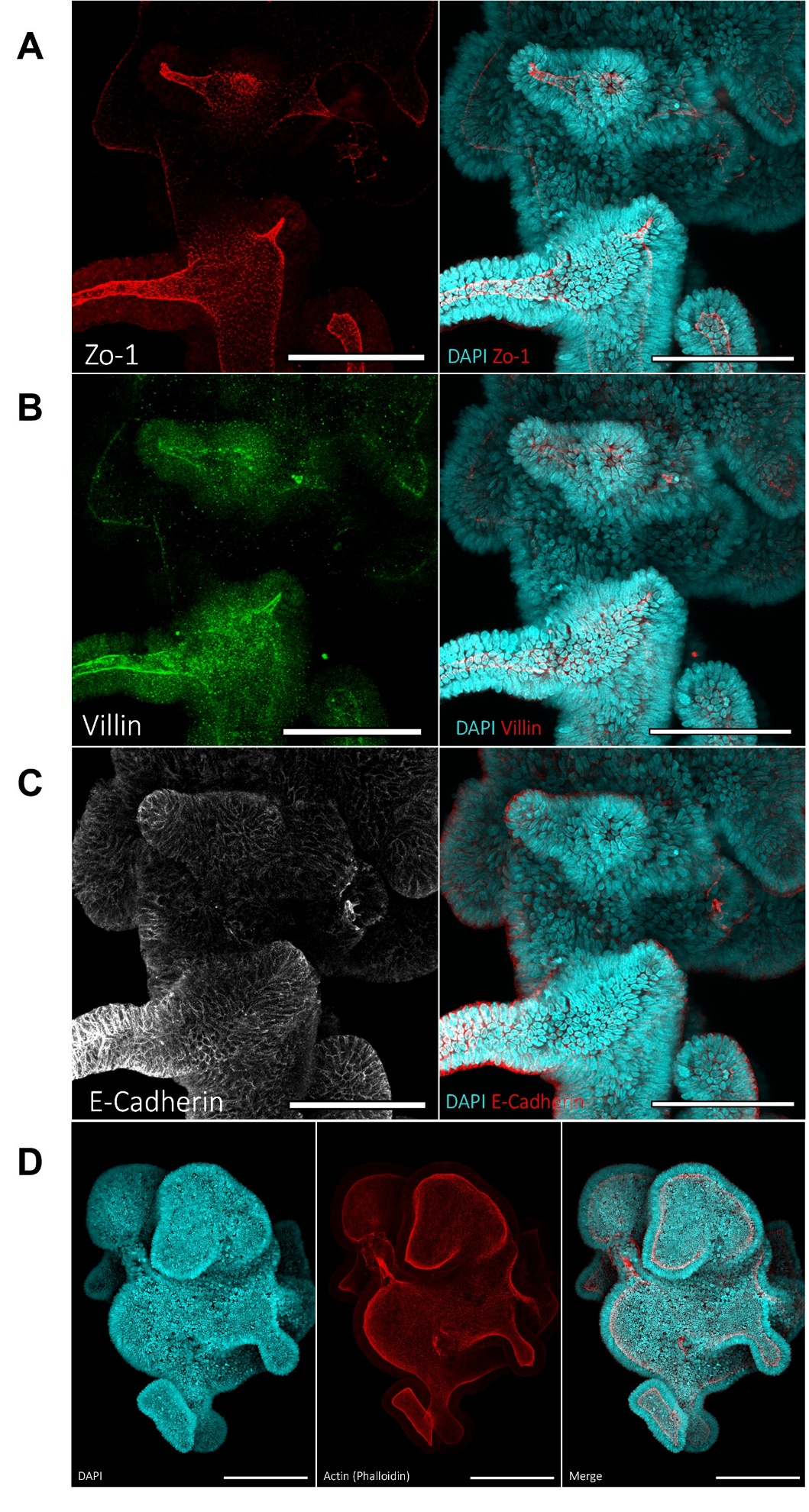

#### Supplementary figure 4. Polarity‑resolving markers demonstrate apical‑in organization in Matrigel‑embedded organoids.

Spatial localization of **(A)** ZO‑1 (red), **(B)** villin (green), and **(C)** E‑cadherin (white) shown individually (left panels) and as a merged overlay with nuclei (cyan, right panels). Marker distribution resolves tight‑junction belts and apical brush border facing the enclosed lumen, consistent with an apical‑in epithelial topology. **(D)** Phalloidin labeling highlights cortical F‑actin at the luminal surface, reinforcing the apical‑in configuration in a large, intact 3D structure. Figures show SpA-derived caecum HIOs collected at day 10 of culture. The samples were fructose–glycerol‑cleared and whole‑mounted on a glass slide. The acquisition was performed on a ZEISS LSM‑700 confocal microscope equipped with a Plan‑Apochromat 20×/0.8 lens. Three-dimensional reconstruction was performed starting from a Z‑stack and 2 x 2-tiled collection in Zeiss ZEN Blue. Scale bar = 100 µm (A, B and C) and 200 µm (D).
